# The influence of visual attention on letter recognition and reading acquisition in Arabic

**DOI:** 10.1101/2024.09.01.610706

**Authors:** Alaa Ghandour, Emmanuel Trouche, Dominique Guillo, Sylviane Valdois

**Affiliations:** Univ. Grenoble Alpes, CNRS, Laboratoire de Psychologie et de NeuroCognition (LPNC), 38000 Grenoble, France; Mohammed VI Polytechnic University, Africa Institute for Research in Economics and Social Sciences, Rabat, Morocco

**Keywords:** Reading acquisition, Arabic Language, Visual Attention Span, Letter Knowledge, Phonological awareness, Cognitive processes, Path Analysis?

## Abstract

The present study sets out to explore the cognitive underpinnings of reading acquisition in Arabic. Previous studies have identified phonological awareness and rapid automatized naming as early predictors. However, the graphic complexity of Arabic letters imposes particular constraints on the visual system, which should mobilize visual attention. To test this hypothesis, 101 Arabic-speaking children who just began their formal reading instruction in Arabic were administered tests of syllable and word reading. Their nonverbal reasoning, vocabulary, phonological awareness, rapid automatized naming and letter knowledge were measured. Their visual attention was estimated through tasks of visual attention span. We found that phonological awareness, visual attention span and letter knowledge were associated with reading outcomes. However, regression analyses showed that the relationship between visual attention span and reading disappeared when letter knowledge was taken-into-account. We used structural equation modeling to examine the direct and indirect effects of visual attention span to reading. Results showed that phonological awareness and letter knowledge were significant and independent predictors of reading while visual attention span contributed only indirectly through its influence on letter knowledge. Our findings suggest that beginning readers rely on visual attention to identify and discriminate visually-complex Arabic letters. In turn, more efficient letter identification in children with higher visual attention facilitates reading acquisition. These findings support the cognitive models of word recognition that include visual attention as a component of the reading system. They open new perspectives for cross-language studies, suggesting that visual attention might contribute differently to reading depending on the orthographic system. They also provide a foundation for innovative teaching methodologies in Arabic language education.

## Introduction

The purpose of this study was to contribute to research into the cognitive skills involved in learning to read. Most previous studies have focused on the Indo-European languages, leading to identifying rapid automatized naming (RAN), phonological awareness (PA) and letter knowledge (LK) as early predictors of learning to read. These same skills were identified across languages, suggesting they might be critical for reading acquisition, whatever language orthography. This is supported by current findings, since RAN, PA and LK have been also identified as contributing to reading development in Chinese and in Semitic languages. However, there is also evidence that the predictive power of these different skills may differ across orthographies [1,2]. The present study focuses on reading acquisition in Arabic. Relatively few studies on reading predictors and reading-related skills have focused on the Arabic language, even though the specific features of this language present particular challenges. In addition to PA, RAN and LK, we will examine the potential impact of visual attention span (VAS). VAS is a measure of multi-element parallel processing that is involved in reading acquisition and developmental dyslexia [3–5]. Its impact on reading has been reported in several languages but evidence for its involvement in Arabic is scarce.

### Arabic Language Specific Features

Arabic is a Semitic language with a rich historical and cultural heritage, distinguished by unique script and linguistic features that contribute to its complexity. Like other Semitic languages, Arabic is written from right to left along a horizontal line. It employs an ABJAD writing system wherein its fundamental script comprises consonants, with optional symbols for denoting short vowels and other morpho-phonemic features of the language [6]. The writing system is substantially more complex in Arabic than in languages using the Latin alphabet [2]. The Arabic alphabet consists of 28 letters, including two semi-vowels that can function as either a consonant or a long vowel depending on context. Written in a cursive style, Arabic orthography features connected letters with no uppercase counterparts. However, the form of Arabic letters varies based on their within-word position (initial, medial, or final). For example, the letter “Kaaf” ﻙ /k/ is written as ﻛـ in the initial position, as ـﻜـ in the medial position, and as ـﻚ in the final position. These distinct letter forms, known as “allographs”, encompass over one hundred variations in the Arabic script. Additionally, despite the cursive nature of the Arabic script, six out of the twenty-eight letters (ﻭ, ﺯ, ﺭ, ﺫ, ﺩ, ﺍ) do not connect to the following letter, resulting in one or more spaces within words. Furthermore, many Arabic letters share similar shapes, differing only in the presence and positioning of dots or points (known as primary diacritics), such as the letters ﺏ /b/, ﺕ /t/, and ﺙ /ɵ/. Short vowels, in Arabic orthography, are represented by secondary diacritics above or below the letter, with three main vowels: /a/ or /fatḥa/, /u/ or /ḍamma/, and /i/ or /kasra/. The absence of a vowel is denoted by or /sukūn/. Certain diacritical marks, such as the /tanwīn/ (or nunation), indicating an indefinite noun through vowel doubling, and the /šadda/ (or gemination), representing consonant doubling, are considered morpho-phonemic [7]. It is important to note that vowelization in Arabic script is optional. Short vowel signs (secondary diacritics) may or may not be included, leading to two versions of the Arabic script: vowelized and unvowelized. The vowelized version is prevalent in classical Arabic texts like the Holy Qur’an, classical poems, and literacy books for young learners. The unvowelized version, devoid of short vowel markings, is used by proficient readers in books, novels, media, etc., and is introduced to children around the fourth grade to gradually familiarize them with reading in its unvowelized form. The two scripts impose different constraints on the cognitive system of reading. The vowelized script is fully transparent while the unvowelized script transcribes only part of the word phonological form [8].

Another fundamental characteristic of the Arabic language is its diglossic nature, wherein it manifests in two distinct forms: the standard variety and the spoken variety. The Modern Standard Arabic (MSA) form adheres to defined rules and grammar, serving as a shared language among all Arabic speakers. It is predominantly used in written form and finds application in formal settings, religious discourse, and media communications [9]. In contrast, the spoken variety is employed in everyday conversations and exhibits geographical variations across regions and even within the same country. This spoken form serves as the primary language of Arabic speakers, acquired naturally through familial interactions. Exposure to MSA typically begins during formal education, often in kindergarten. Notably, a linguistic gap exists between spoken Arabic and MSA, encompassing differences in phonology, lexicon, syntax, and morphosyntax. These disparities position MSA as a second language for young learners [10]. The context of diglossia has received little attention in studies on the cognitive dimensions of reading, even if diglossia is likely to impact the assessment of reading-related language and phonological skills.

Last, Semitic morphology differs from that of European languages due to its unique non-concatenative derivational structure [11]. Arabic morphology relies on a system of discontinuous morphemes known as roots and patterns. The root, typically composed of three consonants, indicates a semantic field, and serves as the foundation for deriving numerous words of the same semantic family. The specific meaning of each word results from the combination of the root with a pattern that corresponds to a set of vowels (and sometimes additional consonants). For instance, from the root KTB, denoting the realm of writing, arise words such as /KaTaBa/ (he wrote), KaaTiB (writer), KiTaaB (book), and maKTaBa (library). There is evidence that the morphological structure of Arabic words has an impact on reading accuracy and comprehension [12–16]. Note that the impact of morphological processing on reading acquisition is beyond the scope of this paper.

### Reading-Related Skills

The process of learning to read is influenced by a variety of linguistic, cognitive, and socio-cultural factors, each of which plays a pivotal role in the development of reading skills. Moreover, the contribution of these factors may vary according to the linguistic characteristics of the language being learned. Depending on the language family (Indo-European languages such as English, Semitic languages such as Arabic, or logographic languages such as Chinese), the interplay between language-specific features and reading predictors underlines the complexity of the reading acquisition process. In the following, we will explore the significance of each of the early predictors of reading proficiency in Indo-European languages, and then review the available evidence for Arabic.

Rapid Automatized Naming (RAN) requires the rapid and accurate naming of arrays of familiar items, such as letters, digits, colors, or objects. Research has consistently shown that RAN is a significant predictor of reading across Indo-European languages [17–22]. Accordingly, deficits in RAN are reported in individuals with developmental dyslexia [23] and RAN weaknesses characterize children at risk for reading difficulties [24]. Although RAN remains a significant predictor of reading in transparent orthographies [25], it may have stronger influence on reading fluency in opaque languages [21]. RAN also contributes to reading skills in Arabic. Performance in RAN-letter or RAN-digit is a significant predictor of Arabic reading [26–27] but RAN-object also contributes to reading speed [28]. Furthermore, deficits in RAN correlate with reading difficulties in Arabic-speaking children, highlighting its importance in identifying individuals in need of additional literacy support [29]. Therefore, RAN emerges as a critical predictor of reading development in Arabic.

Phonological Awareness (PA), the ability to recognize and manipulate the phonological units of spoken language, is a key predictor of reading achievement in all languages. Strong PA skills facilitate accurate decoding and fluent reading, contributing to both early reading acquisition and later reading comprehension [30–32]. PA at the syllable and rhyme level is a less effective predictor than phoneme awareness [33]. The strength of the relationship between PA and reading may also vary according to language transparency [32,34–36]. Phoneme awareness develops earlier in transparent than in opaque orthographies [37] and syllable awareness may play a stronger role in transparent languages with simple syllable structure [34]. Similar to other languages, phonological awareness is a strong predictor of reading proficiency in Arabic [38–41]. PA is a significant predictor of Arabic reading, independent of RAN [26,38], and poorer PA skills have been reported in at-risk children and Arabic dyslexic readers [42,39]. Good PA in preschool Arabic children is also critical for literacy development [43]. According to Saiegh-Haddad et al. [44], the phonological distance between MSA and spoken Arabic can have a noteworthy impact on the quality of phonological representations across all academic levels. This linguistic duality poses unique challenges for the development of phonological awareness, so the diglossia context should be considered when designing phonological awareness tasks [45,46].

Letter knowledge (LK), defined as the ability to recognize letter shapes and associate them with the corresponding name or sound, also contributes to reading achievement in Indo-European languages [47]. Letter Knowledge is one of the strongest predictors of early reading skills, in both transparent and opaque languages [25,48–51]. Children with higher preschool letter knowledge develop more efficient word recognition skills, with subsequent positive effects on reading comprehension [52]. Letter knowledge also contributes to the development of phoneme awareness and decoding skills [53,54]. Assuming that word processing depends on efficiency in letter identification [55], letter knowledge, and more specifically the ability to recognize and distinguish letters based on their visual features, should contribute to reading in all languages. In Arabic, letter identification is known to be challenging for beginning (and even more proficient) readers [56,57]. In particular, a number of studies have highlighted the difficulty of recognizing and discriminating between Arabic letters, suggesting that letter recognition may be more demanding on visual attention, thus resulting in slower reading [58,59]. However, we lack direct evidence that letter knowledge predicts reading in Arabic and, to our knowledge, no study has investigated the potential unique contribution of letter knowledge to reading, beyond that of PA or RAN.

A last factor influencing reading acquisition is the visual attention span [60]. Visual Attention Span (VAS) is a measure of multi-element parallel processing in the visual modality that affects reading ability by determining the number of letters that can be processed simultaneously [60– 62]. VAS contributes to decoding, word recognition and fluency, beyond PA and RAN [3,63,64]. It is an early predictor of later reading skills [65], so that VAS is impaired in children with reading difficulties and developmental dyslexia [4,61,66]. Although an effect of VAS on reading acquisition has been reported regardless of language transparency, a recent study by Liu et al. [5] showed that the VAS deficit in dyslexic individuals was greater in opaque than in transparent languages. The few studies, that have investigated the contribution of VAS to reading in Arabic, have produced inconsistent results. Awadh et al. [67] investigated whether VAS predicted text reading fluency in expert readers of Arabic, French and Spanish. They reported no effect of VAS on (non-vowelized) text reading in Arabic. In contrast, a relationship between VAS and text reading was reported by Lallier et al. [68] in Grade 4 Arabic children but only for the participants who were more proficient in reading non-vowelized scripts. Finally, Awadh et al. [69] investigated both VAS and PA skills as predictors of reading fluency and comprehension in Grade 4 and Grade 5 native Arabic readers. They showed that VAS uniquely contributed not only to word and pseudoword reading but also to text fluency and comprehension. These inconsistent findings highlight the need for further research into the VAS-reading relationship in the Arabic language.

In summary, RAN, PA, LK and VAS are all recognized as early and independent predictors of reading achievement in Indo-European languages. The respective roles of PA and RAN have also been investigated in Arabic, with similar evidence for the unique contribution of each of these two skills to reading. The potential contribution of LK has not been investigated independently of PA and RAN, and the potential role of VAS as an early and independent predictor of reading in Arabic is still inconclusive.

### The present study

The aim of the present study was to assess whether PA, RAN, LK and VAS each contribute independently to reading skills in Arabic beginning readers. To avoid strong relationships between the different predictors, RAN, LK and VAS were assessed using different stimuli. LK assessment required the use of Arabic letters but both RAN and VAS tap into processes that can be measured using other types of stimuli. RAN can be measured using either alphanumeric or non-alphanumeric items to tap rapid access to phonological labels, fast visuo-verbal matching and/or processing speed. Participants were administered a RAN-Object task, as a relationship between this version of the RAN task and reading in Arabic has been previously reported [28]. VAS is measured using tasks that require the processing of briefly presented strings of visual stimuli. Similar performance is obtained regardless of stimulus type (alphanumeric or not), and performance is interpreted as reflecting the amount of visual attention devoted to the simultaneous processing of multiple elements [60]. Taking advantage of the bilingualism of the participants, the VAS report tasks were designed using Latin letters. The use of Arabic letters for the assessment of LK, objects for RAN and Latin letters for VAS would allow each basic skill to be more specifically tapped.

In line with previous behavioral data and with reference to current knowledge about reading acquisition [33], higher PA at the start of literacy instruction should facilitate the development of the alphabetic principle and thus predict higher word and syllable reading in vowelized script. Similarly, RAN should account for a significant amount of variance in Arabic reading, over and above PA. Letter knowledge is critical in the early stages of reading development across languages and may contribute even more to reading in Arabic due to the visual complexity and high similarity between Arabic letters. Based on previous evidence from alphabetic languages [47], the contribution of LK to reading was expected to be independent of that of PA and RAN. A key originality of the present study was to assess VAS ability as a potential additional predictor of reading in Arabic, beyond PA, RAN and LK. Based on previous evidence from Indo-European languages [4,64], a significant relationship between VAS and reading was expected in Arabic, independent of PA and RAN. However, empirical findings suggest that the processing of Arabic letters may be particularly demanding on visual attention [59]. The involvement of visual attention (here indexed by the VAS measure) in the recognition of individual Arabic letter would predict a significant relationship between VAS and LK, which might affect the VAS-reading relationship. Then, either VAS would have some direct residual effect on reading while controlling for LK or the VAS-reading relationship would be fully mediated by LK, so that VAS would only indirectly affect reading skills through LK.

Reading skills were assessed through tasks of syllable and word reading. Evidence from Indo-European languages suggests a differential involvement of each of the PA, RAN, LK and VAS skills in word and pseudoword reading [70,71]. Therefore, we will assess the predictive power of these different skills independently for nonsense syllables and for real words.

## Materials and Methods

### Participants

One hundred and thirty-four bilingual Lebanese first graders were recruited from four private schools with mild to low socio-economic levels. They had normal hearing and either normal or corrected-to-normal visual acuity. All participants had completed three years of kindergarten and began formal reading instruction in grade 1, with letter teaching starting in KG3. They had Arabic as primary language and English as second language or as language of instruction. All children used Lebanese Arabic at home and were exposed to both the English and the Arabic language at school from kindergarten. The children were tested during the second trimester of the academic year (February – March 2023), after four months of formal Arabic reading instruction. The study was conducted in accordance with the ethical principles expressed in the Declaration of Helsinki. Ethics approval for the study (PASEM Project: E.T. as PI) was granted by the Ethic Committee of the Africa Institute for Research in Economics and Social Sciences (ECAIRESS-002-2024). Legal responsibility for the children during school hours was assumed by the school directors, who provided consent for the students to participate and sought consent from all parents for their child’s involvement. Additionally, verbal consent was obtained from each child at the beginning of every test session, and they were reassured that they could stop participating whenever they wished.

Due to the absence of children during at least one day of data collection, we were unable to collect data for all tasks on all children: Eighteen data points were missing for the reading tasks, twenty-three for the visual attention span tasks and ten for the phonological awareness tasks, resulting in the exclusion of 32 children (24%) who had missing data in at least one of these skills. In addition, we excluded one extreme outlier, very likely due to a measurement error during the reading task (120 syllables correctly read per minute). Further analysis relies on a final sample of 101 first graders (58 females) who were 7-year old on average (mean age=6 years11 months; SD=4.1 months).

### Measures

Most of the tasks we used were custom-designed due to the absence of standardized task for assessment in beginning Arabic readers.

### Non-verbal Reasoning Test

The Raven Colored Progressive Matrices (RCPM) test assessed participants’ abstract reasoning abilities as a measure of fluid intelligence [72]. Participants were presented with three series of 12 colored patterns arranged in matrices and asked to select the missing pattern from a set of options provided below each matrix. In the absence of normative data for the Lebanese population, we used the children’s raw scores (maximum score = 36).

### Vocabulary Knowledge

Participants completed an object naming test derived from the ELO-L, a language screening test for 3 to 8-year-old Lebanese children [73]. The test consisted of 35 pictures, each presented individually. Participants were encouraged to respond in standard Arabic but responses in spoken Arabic were accepted if they were unfamiliar with the standard Arabic label. The score was calculated as the total number of correct responses, regardless of the language register used (maximum score = 35).

### Rapid Automatized Naming (RAN)

Participants completed a custom-designed RAN task, in which they had to quickly name a series of familiar objects. The task consisted of 5 different objects that were repeated 5 times each and presented in a random order. All words were monosyllabic high frequency words (according to the ALEF frequency database for Grades 1 and 2 [74]) that showed minimal variation between MSA and spoken Lebanese Arabic: bear (/dub/ in MSA - /dib/ in Lebanese), rooster (/diik/-/diik/), hand (/yad/-/iid/), elephant (/fiil/-/fiil/), and house (/bayt/-/beet/). First, participants were checked to ensure that they could recognize and name each picture accurately. Then, they were asked to name all 25 pictures (arranged in a 5 x 5 array) horizontally from right to left as quickly as possible. The experimenter recorded the total time taken to complete the task (expressed in seconds).

### Phonological Awareness (PA)

Participants’ phonological awareness was assessed using three tasks of Syllable Segmentation (SS), Initial Syllable Deletion (ISD), and Initial Phoneme Deletion (IPD). The Syllable Segmentation task was specifically designed for this study, while we used the deletion tasks from the BELEA battery [75]. All tasks were administered in MSA, using items with minimal differences from spoken Lebanese Arabic. Each task consisted of 8 items, preceded by 3 to 5 practice items. Both scores and time were recorded.

In the Syllable Segmentation task, participants had to segment 5 2-syllable-words with simple CV and CVV syllables (e.g., /saa-’a/ meaning "hour" or "clock") and 3 3-syllable-words with, at least, one complex CVC syllable (e.g., /laa-’i-bun/ meaning "player").

In the Syllable Deletion task, children were asked to mentally delete the first syllable of familiar 2- or 3-syllable words and say what remained. The syllables to be dropped were either simple CV and CVV syllables (e.g., /baa-ri-dun/ meaning "cold") or complex CVC syllables (e.g., /Sham-sun/ meaning "sun").

In the Phoneme Deletion task, participants had to omit the initial phoneme from words consisting of 2-to-3 syllables and 5-to-7 phonemes. To reduce memory load, all words shared their last 2 phonemes (corresponding to the nunation /un/ that indicates that they were syntactically indefinite). In 5 of the 8 items, the first phoneme was omitted from a long syllable where the vowel was represented by a whole letter (e.g., /fii-lun/ meaning "elephant"), and in the remaining 3 items, it was omitted from a short syllable where the vowel was represented by a diacritical mark (e.g., /qal-bun/ meaning "heart").

### Letter Knowledge (LK)

The 28 Arabic letters were presented individually in random order. The children were instructed to name each letter as quickly and accurately as possible. They were asked to say the name of the letter. However, some participants responded with the letter sound combined with the vowel /a/ (e.g., /da/ for the letter /daal/; /sa/ for the letter /siin/). As some children may have been taught letter names while others may have been taught letter sounds, all these types of responses were scored as correct similar to what has been done by Tibi et al. **[76]** (maximum score = 28).

### Visual Attention Span (VAS) and Single Letter Identification Threshold

Participants’ visual attention span was assessed using global and partial letter report tasks [60]. These tasks used strings of four Latin letters (e.g., R H S D) constructed from ten consonants (B, P, T, F, L, M, D, S, R, H). Letters were presented in uppercase black on a white background, with no repeated letters or real-word patterns within a string. All tasks were administered using the E-Prime software. On each trial, a central fixation point appeared for 1000ms, immediately followed by the letter string presented for 200ms. In the global report task, 20 4-letter strings were presented successively, and participants had to report as many letters as possible in any order at the string offset. In the partial report task, the briefly presented 4-letter string was immediately followed by a cue indicating the position of the letter to be reported. Forty trials were presented in succession (10 targets per position). Feedback was given during training but withheld during experimental trials. One point was scored for each correctly reported letter, with a maximum score of 80 for the global report and 40 for the partial report tasks.

Further, a single-letter identification threshold task was administered to control for single Latin letter processing speed. Each of the 10 consonants used in the report tasks was presented in isolation for randomly varying durations from 33ms to 101ms (in 16ms increments), followed by a mask to erase the information from iconic memory. One point was awarded for each letter accurately named. Following Bosse & Valdois [63], the total score was calculated as the sum of the weighted scores at each presentation duration (5 times the score at 33ms + 4 times at 50ms + 3 times at 67ms, 2 times at 84ms, and once at 101ms; leading to a maximal score of 150).

## Reading

Three custom-made lists of 12 items each were designed to assess reading fluency for nonsense syllables, monosyllabic words, and polysyllabic words. All items were written in a fully transparent, vowelized script and were from 2-to-4 letter long. They were chosen to show minimal differences between spoken Lebanese and standard Arabic. The words were presented in columns, and each list was read separately. The list of nonsense syllables (one- or two-letter long) included 5 short syllables (CV) where the vowel is represented by a diacritical mark (e.g., /si/ ﺱِ), with consonants varying in frequency and no rare consonants used [77], and 7 long syllables (CVV) in which the vowel is represented by a whole letter (e.g., /fuu/ ﻓﻮ), with 5 of the 7 syllables considered to have a high level of discriminability and 2 having a high level of difficulty [78]. The second list consisted of 1-syllable complex words that were two-to-three-letter long and included frequent patterns (ALEF database, [74]), namely either a CVC (e.g., /Ɂaχ/ meaning "brother"), CVVC (e.g., /ҁiid/ meaning "holiday"), or CVCC (e.g., /Ṣayf/ meaning "summer") structure. The third list consisted of 2-to-3-syllable simple words that were three-to-four-letter long and included frequent patterns (ALEF database, [74]), namely CVV-CVV (e.g., /raa-mii/ which is a proper name), CVV-CV (e.g., /naa-ma/ meaning "he slept"), CV-CV-CV (e.g., /ka-ta-ba/ meaning "he wrote" and /ʃa-ri-ba/ meaning "he drank"). Participants were instructed to read each list as quickly and accurately as possible. Scores and time (in seconds) were recorded for each list, with a maximum score of 12.

## Data Collection

Data collection was carried out by experimenters, all of whom were trained speech and language therapists or trained undergraduate speech and language therapy students. The tests were administered individually in a quiet room or hall at school, with the order of testing randomized to avoid bias due to fatigue effects. Each child was tested in 2-3 sessions, each lasting 35 to 50 minutes. Following the administration of the various tasks, scoring was carefully completed by the experimenters and cross-checked twice — first by a research assistant and then by the lead researcher of the study — to avoid input errors. Reliability estimates according to McDonald’s Omega coefficients [79] for the different measures are presented in Table 1.

**Table 1.** Descriptive statistics and McDonald’s Omega reliability coefficients (ω) for all variables. Mean (M), Standard deviation (SD), Median (Mdn), Minimum (Min.), Maximum (Max.), Skewness (Skew.), Kurtosis (Kurt.), data transformation applied in further analysis (Transf.)

| | *M* | *SD* | *Mdn* | *Min.* | *Max.* | *Skew.* | *Kurt.* | *Transf.* | $\omega$ |
| --- | --- | --- | --- | --- | --- | --- | --- | --- | --- |
| Raven | 17.18 | 4.24 | 17.00 | 6.00 | 31.00 | 0.48 | 1.18 | none | .76 |
| Vocabulary | 12.37 | 5.38 | 13.00 | 1.00 | 25.00 | 0.07 | -0.66 | none | .85 |
| RAN | 39.39 | 17.93 | 34.00 | 19.00 | 118.00 | 2.20 | 5.51 | $\frac{1}{x}$ | NA |
| LK | 20.19 | 5.82 | 22.00 | 6.00 | 28.00 | -0.64 | -0.52 | none | .91 |
| PA | 5.34 | 3.27 | 4.82 | 0.35 | 16.98 | 1.17 | 1.42 | $\sqrt{x}$ | .85 |
| • Syllable Segmentation | 9.57 | 4.80 | 9.06 | 0.44 | 20.87 | 0.20 | -0.63 | NA | .84 |
| • Syllable Deletion | 3.86 | 3.28 | 3.00 | 0.00 | 15.56 | 1.26 | 1.25 | NA | .85 |
| VAS | 45.82 | 13.11 | 45.00 | 14.00 | 71.50 | -0.09 | -0.50 | none | NA |
| • Partial Report | 24.89 | 8.20 | 25.00 | 3.00 | 38.00 | -0.49 | -0.35 | NA | NA |
| • Global Report | 41.86 | 12.63 | 40.00 | 12.00 | 73.00 | 0.33 | -0.40 | NA | NA |
| Letter Threshold | 90.04 | 34.58 | 96.00 | 6.00 | 150.00 | -0.22 | -1.01 | none | NA |
| Syllable Reading | 11.33 | 13.60 | 6.00 | 0.00 | 80.00 | 2.30 | 6.60 | $\sqrt{x}$ | .91 |
| Word Reading | 4.45 | 5.76 | 2.75 | 0.00 | 28.80 | 2.31 | 5.56 | $\sqrt{x}$ | .95 |
| • Monosyllable Words | 3.75 | 5.39 | 2.09 | 0.00 | 24.44 | 2.57 | 6.39 | NA | .89 |
| • Multisyllable Words | 5.67 | 7.06 | 2.50 | 0.00 | 36.00 | 2.07 | 4.61 | NA | .92 |

Note: RAN= rapid automatized naming; LK=letter knowledge; PA= phonological awareness; VAS= Visual attention span.

## Results

### Descriptive Statistics

We observed a massive floor effect on the Phoneme Deletion task, with 68% of the children getting a score of 0 over 8, so that the measure was removed from further analysis. Table 1 presents the descriptive statistics for all the other variables, and for three composite measures of PA, VAS and Word Reading. The composite score for PA was calculated by averaging performance on the Syllable Segmentation and Syllable Deletion tasks. The composite score for Word Reading corresponds to the average of the scores on the mono- and multi-syllable words. Following Ginestet et al. [80], the composite VAS score was calculated using the following formula:

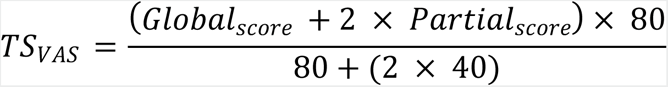

Before computing the composite scores of PA, VAS and Word Reading, we checked that the scores of the two tasks used to create each of them were correlated. The correlations between Syllable Segmentation and Syllable Deletion [*r*(99) = .36, *p* < .001], Partial and Global Report [*r*(99) = .62, *p* < .001] and, Monosyllable and Multisyllable Words [*r*(99) = .87, *p* < .001] were significant. These correlations, together with the high Omega coefficients of internal consistency when all items are aggregated (ω_PA_= .85; ω_VAS_=; ω_Word Reading_= .95), support our decision to use these constructs in further analyses.

As shown in Table 1, the McDonald’s Omega of all the variables, including our three constructs, were above .76, indicating good internal consistency of our measurement tools. The distributions of most variables (Raven, Vocabulary, LK, Letter Threshold and VAS) were close to normal, with skewness values ranging from -.64 to .48 and kurtosis from -1.01 to 1.18. The distribution was moderately skewed for PA and highly skewed and peaked for the three measures of RAN, Syllable Reading and Word Reading. To increase their symmetry and reduce their kurtosis, square root transformations were applied to all these variables except RAN, for which an inverse transformation was applied. After data transformation, all the skewness and kurtosis values ranged from -.87 to 1.01 and 1.01 to 1.18 respectively (see Table S1 in supplementary material).

### Correlation Analyses

Table 2 shows the simple and partial correlation coefficients (after controlling for Raven) between all our main variables.

**Table 2.** Pearson Correlations (above the diagonal) and partial correlations (below the diagonal) after control of Raven.

|  | 2 | 3 | 4 | 5 | 6 | 7 | 8 | 9 |
| --- | --- | --- | --- | --- | --- | --- | --- | --- |
| 1 - Raven | .14 | .25 | .14 | .27 | .15 | .38** | .33 | .31 |
| 2 - Vocabulary | - | .28 | .19 | .01 | .12 | .09 | .19 | .19 |
| 3 - RAN | .25 | - | .17 | .32 | .06 | .26 | .35* | .35* |
| 4 - LK | .18 | .14 | - | .26 | .27 | .58*** | .72*** | .66*** |
| 5 - PA | -.03 | .27 | .24 | - | .31 | .43*** | .50*** | .58*** |
| 6 - Letter Threshold | .11 | .03 | .25 | .28 | - | .40** | .29 | .25 |
| 7 - VAS | .04 | .18 | .57*** | .37* | .37** | - | .53*** | .58*** |
| 8 - Syllable Reading | .16 | .29 | .72*** | .45*** | .26 | .46*** | - | .84*** |
| 9 - Word Reading | .15 | .30 | .65*** | .54*** | .22 | .52*** | .82*** | - |

*Note.* \* $p < .05$ ; \*\* $p < .01$ ; \*\*\* $p < .001$ ; $p$ -values are adjusted using Bonferroni correction for 64 tests

The results show that the predictive variables of PA, LK and VAS were highly correlated with the two dependent measures of syllable and word reading. RAN was significantly correlated with both reading measures but only when Raven was not controlled for. In contrast, Vocabulary was not significantly related to any measure and was therefore excluded as predictor from further analyses. The highest correlations were observed between the two word and syllable reading tasks, and between each of them and LK. However, significant correlations were also found between some of the predictive variables. In particular, VAS correlated moderately with LK [*r*(99) = .57, *p* < .001] and to a lesser extent with PA [*r*(99) = .37, *p* < .05].

### Regressions Analyses and Structural Equation Modeling

Simple regression analyses were performed with either Syllable Reading or Word Reading as the dependent variables and PA, RAN, VAS, Raven, Vocabulary and Letter Threshold as independent variables. The analyses revealed that RAN was not a significant predictor of reading fluency (for Syllable and Word Reading respectively, *t* = 1.14, *p* = .17, *R_p_ ^2^* = .02 and *t* = 1.05, *p* = .30, *R_p_ ^2^* = .01). In fact, only PA and VAS were significant predictors of reading skills (for Syllable and Word Reading respectively, PA: *t* = 3.15, *p* = .002, *R_p_ ^2^* = .10, and *t* = 4.64, *p* < .001, *R_p_ ^2^* = .19; VAS: *t* = 3.24, *p* = .002, *R_p_ ^2^* = .10, and *t* = 4.24, *p* < .001, *R_p_ ^2^* = .16, see Tables S2 and S3 in supplementary material).

However, when adding LK as an additional independent variable, PA became an even better predictor of reading (Syllables: *t* = 3.83, *p* < .001, *R_p_ ^2^* = .14; Words: *t* = 5.26, *p* < .001, *R_p_ ^2^* = .23), while the predictive effects of VAS completely disappeared (Syllables: *t* = -0.65, *p* = .51, *R_p_^2^* < .01; Words: *t* = 0.89, *p* = .28, *R_p_^2^* = .01). RAN improved its predictive effect although not to the point of reaching significance level (Syllables: *t* = 1.87, *p* = .07, *R_p_^2^* = .04; Words: *t* = 1.27, *p* = .21, *R_p_^2^* = .02). Remarkably, LK became the best predictor of Syllable and Word Reading (respectively*, t* = 8.35, *p* < .001, *R_p_^2^* = .43, and *t* = 6.24, *p* < .001, *R_p_^2^* = .30). These results, along with the significant correlation (*r* = 0.57) between LK and VAS, suggest that VAS may only contribute to reading performance indirectly, through LK.

To directly test this hypothesis, we performed two structural equation models, similar to the previous simple regressions except that they included both the direct and indirect (through LK) effects of VAS on reading performance. As shown in Fig 1 and 2, the models explained respectively 67% and 65% of Syllable Reading and Word Reading respectively (see Tables S4 and S5 in supplementary material for more information about the two structural models). The two models enabled us to confirm our hypothesis, *i.e.,* VAS did not have any direct effect on reading (Syllables: *β* = -.06, *p* = .50, *z* = -0.68; Words: *β* = .10, *p* = .25, *z* = 1.14) but only an indirect one through LK (Syllables: *β* = .37, *p* < .001, *z* = 5.58; Words: *β* = .28, *p* < .001, *z* = 4.88). VAS explained 34% of LK’s variance, and LK explained respectively 43% and 30% of Syllable Reading and Word Reading variance.

**Fig 1.**
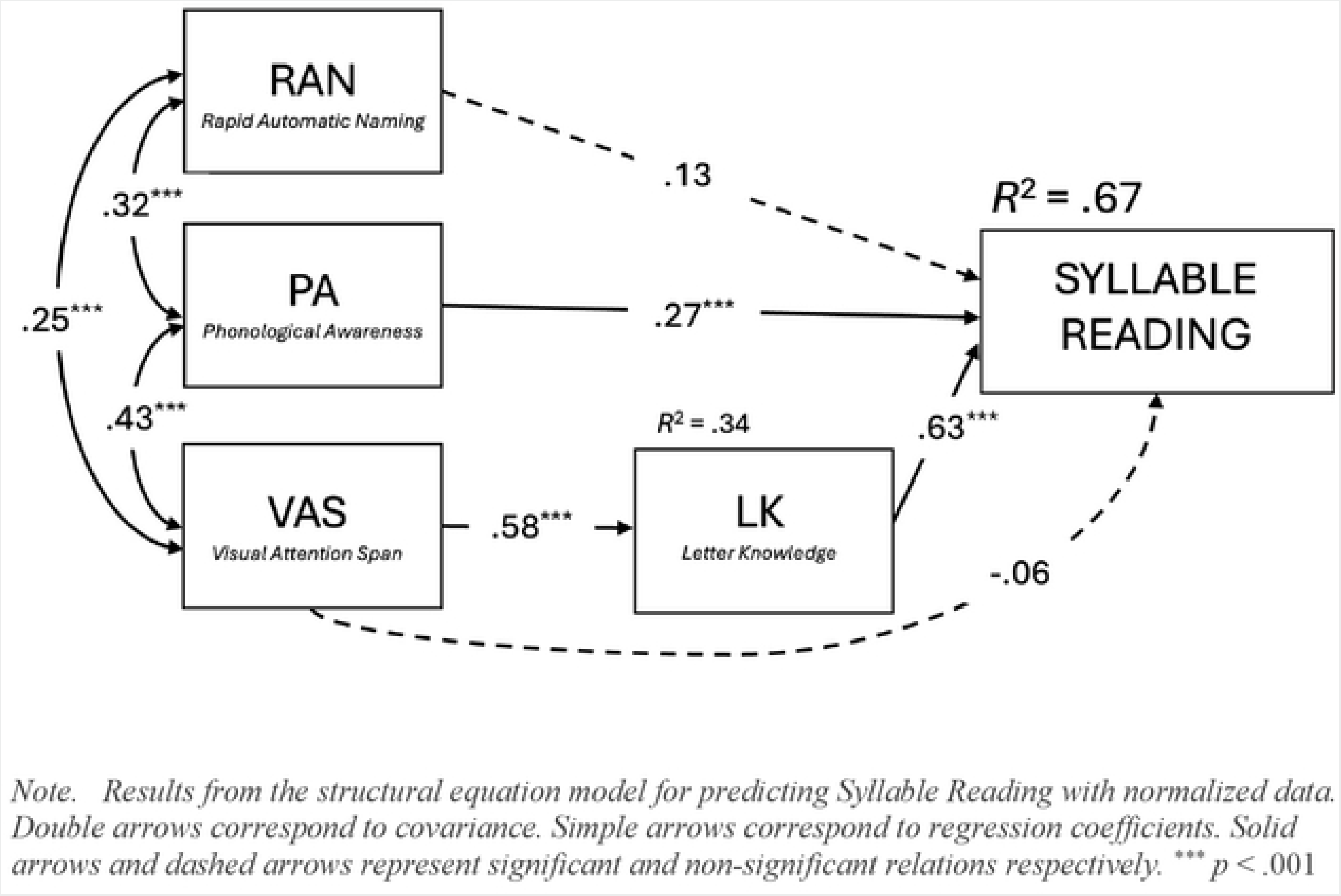
**Predictors of Syllable Reading after controlling for Raven, Vocabulary and Letter Threshold.**

**Fig 2.**
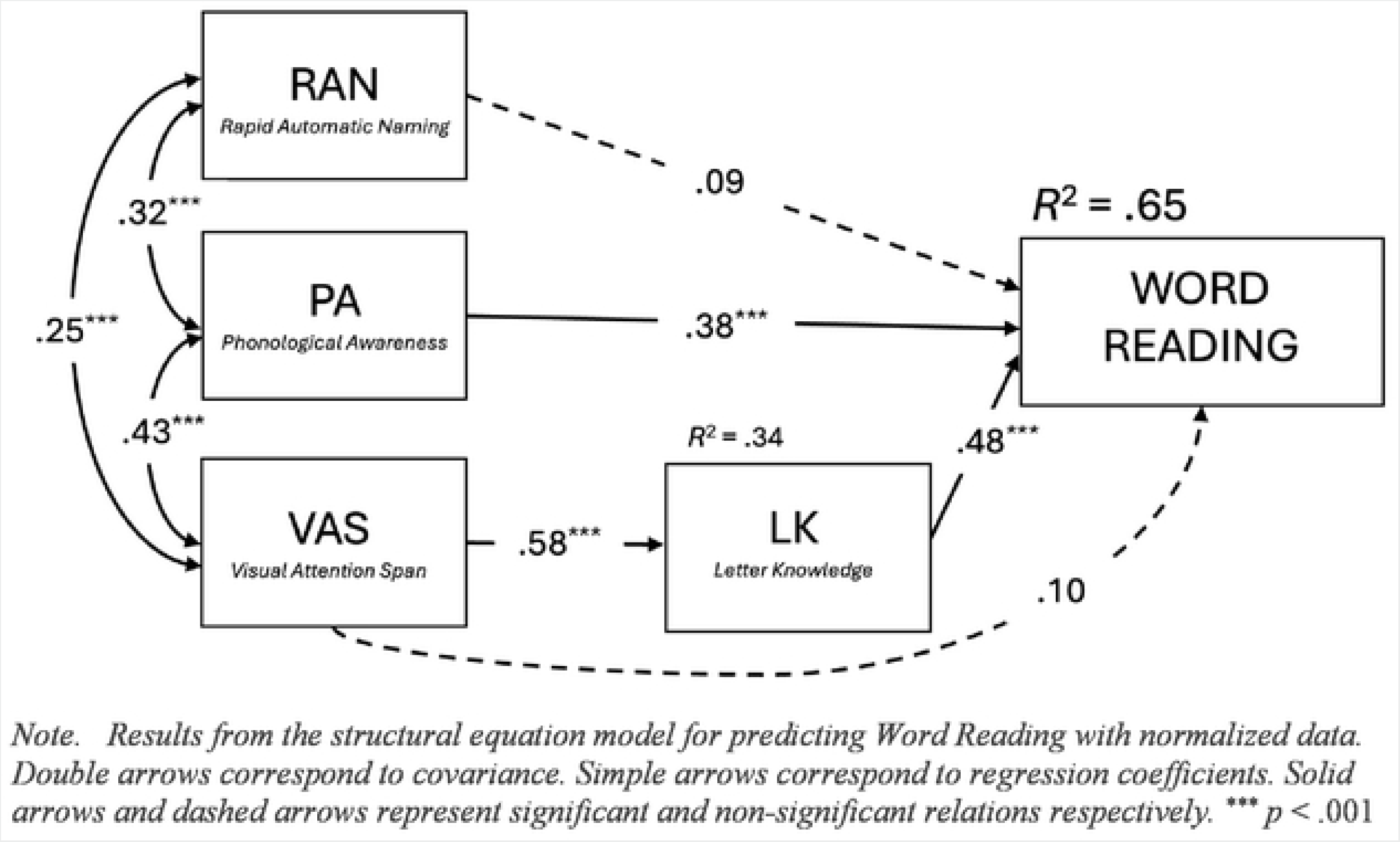
**Predictors of Word Reading after controlling for Raven, Vocabulary and Letter Threshold.**

## Discussion

The goal of this study was to identify the concurrent predictors of reading fluency in beginning Arabic readers. To achieve this, we assessed both phonological and visual processing skills, with a particular emphasis on letter knowledge and visual attentional processing. Consistent with previous studies [38,41,69,81–86], correlation analyses confirmed that children who performed best in syllable and word reading also exhibited higher performance in phonological awareness (PA). However, the relationship between reading and Rapid Automatized Naming (RAN) of objects was not significant after controlling for general nonverbal ability. This finding contrasts with a number of previous studies [41,42,87–89] that reported a significant relationship between RAN and different reading measures in older Arabic-speaking children (Grades 3 to 5). Similarly, Asaad & Eviatar [90] found significant RAN-reading relationships in Grade 1 Arabic readers, but their study focused on the link between RAN, measured with Arabic letters, and text reading speed.

Several studies have consistently found positive correlations between RAN and various reading tasks across age ranges from KG3 to Grade 6 [26,38,41,42,88,91–93]. These studies have examined different types of stimuli, including digits, letters, objects, colors, and shapes. However, the specific choice of items is rarely detailed, with little consideration given to factors such as diglossia, the number of syllables, or item frequency. Additionally, some studies report correlations between reading and composite RAN factors that combine object and digit or object and letter naming. Consequently, due to the inconsistency of stimulus types and characteristics, the role of RAN remains unclear across the literature.

Like in Indo-European languages [47], we show that letter knowledge (LK) is the skill most strongly associated with reading proficiency in our sample of beginning Arabic readers. Some studies have highlighted the importance of LK in the acquisition of reading skills in Arabic, as it serves as a foundational element for phonological processing and word recognition [78,94]. For instance, children with higher letter recognition abilities demonstrate better reading accuracy and fluency, indicating that a strong grasp of the Arabic script is crucial for developing reading skills [90,95].

Interestingly, children whose reading was more efficient also exhibited a higher Visual Attention Span (VAS). While a significant VAS-reading relationship has not been demonstrated in Arabic-speaking adults when reading non-vowelized text [67], it was reported in children from Grades 4 to 5 when reading vowelized words [69]. Correlation analyses also revealed that VAS was related to LK, underscoring the interconnectedness of visual processing skills and letter recognition in reading acquisition. Children with better visual attention abilities tend to have higher letter knowledge, possibly due to their enhanced ability to focus on and discriminate between visually similar Arabic letters, which often share common features and can be easily confused [78,77,96]. This interrelation highlights the importance of integrating both visual and phonological training in educational strategies to support reading development in Arabic.

We further showed that both VAS and PA were significant predictors of reading fluency when LK was not introduced as predictor in regression analyses. However, the predictive power of VAS disappeared when LK was an additional predictor. Last, the predictive power of each variable was studied while considering the direct and indirect effects of VAS to reading.

First, our results confirmed the importance of PA in reading acquisition for the Arabic language [26,97,98]. They suggest that PA is a better predictor of reading than RAN, which had already been highlighted in Arabic beginning readers [38,97]. As emphasized above, the absence of significant RAN-reading relationship in the present study could be due to the use of objects rather than letters or digits. However, the strength of the RAN-reading relationship may also vary depending on whether the influence of visual attention is considered or not. Indeed, despite significant RAN-reading-speed correlations, Layes et al. [87] reported no significant contribution of RAN to reading, beyond that of visual attention (measured through a cancellation task). Further investigation is required to determine whether RAN (measured using letters or digits) would explain additional variance in Arabic reading after controlling for both PA and visual attention.

Our main purpose was to examine whether VAS was a significant and direct predictor of reading skills, independently of PA and LK (after controlling for RAN and for general vocabulary and IQ ability), or whether it contributed only indirectly to reading through LK. First, we showed that LK was a significant unique predictor of reading in Arabic, and that its predictive power was higher than that of PA (or RAN). These findings are fully consistent with previous studies in Indo-European languages, most of which concluded that LK was the strongest concurrent and longitudinal predictor of reading in beginning readers [47,99,100]. The pivotal role of letter knowledge in reading acquisition has recently been confirmed by the results of a training study which used an artificial orthography learning paradigm [53]. The participants were first trained to learn a set of unfamiliar symbols through naming and copying tasks. Then, they were exposed to artificial words made up of the same symbols while listening their pronunciation. After the learning phase, the children who had learned the letters more efficiently showed higher reading accuracy on trained words and higher ability to identify word spellings among distractors. These findings suggest that letter knowledge causally relates to word reading.

Second and more importantly, our findings provide first evidence that visual attention contributes to the processing of individual Arabic letters. It is well documented that the higher visual complexity of Arabic than Latin letters translates in less accurate and slower identification [2,55,56,59]. This led to hypothesize that the processing of Arabic letters might require more visual attention [13,58,85]. Our results demonstrate that visual attention is involved in Arabic letter identification. Assuming that a letter is recognized whenever enough of its component features have been identified [55,100] and that VAS reflects the amount of attention that is deployed for the simultaneous processing of multiple visual elements [60], the link that we highlight between VAS and LK suggests that children who have greater visual attentional resources would process more visual features simultaneously, thus being able to recognize individual Arabic letters more accurately (see [102], for a similar account). Although it is thought that the same mechanisms are involved in all languages, no similar contribution of VAS to letter recognition was reported for Latin letters at the end of kindergarten [65], suggesting lower visual attention involvement during the processing of visually-less-complex Latin letters. These findings could be related to the results of a recent study that showed greater recruitment of the right and left superior parietal lobules during the copy of Arabic than Latin letters [103]. Interestingly, these parietal regions that belong to the dorsal attentional network have been identified as the neural underpinnings of VAS [104 –107].

Last but not least, our findings show that, despite strong VAS-reading correlations, visual attention is not a direct predictor of reading performance when LK is considered as an additional predictor. This contrasts with previous evidence for a direct longitudinal contribution of VAS to reading fluency in French, after controlling for PA and LK [65]. VAS has also been reported as a significant, direct and independent, predictor of reading in many other studies but most of these were carried out on more skilled readers and did not include letter knowledge as a concurrent predictor [3–5].

To better understand this apparent inconsistency, we need to draw on theoretical models of word recognition that include visual attention as a critical component [102,108,109]. It is widely assumed that letter identification within words is more or less efficient depending on their within-string position (due to the acuity and crowding effects) and on the number of features they share with the other letters of the alphabet [101]. In the framework of the BRAID model [108,109,110,111], visual attention facilitates letter recognition, by counterbalancing the deleterious effects of acuity, crowding and cross-letter visual similarity. Thus, more attention is needed to accurately process the component features of low-discriminable letters. In the case of Arabic letters that are characterized by high graphic complexity and low discriminability, a large amount of attention would be allocated to each individual letter for their accurate identification, thus limiting the number of letters that can be simultaneously processed. Thus, in beginning readers who are not yet familiar with Arabic letters, syllables and words might be processed letter-by-letter, leading reading performance to be mainly dependent on the efficiency of each successive letter identification. On the other hand, because of their lesser complexity and higher discriminability, Latin letters might be less demanding in visual attention, thus allowing more letters to be simultaneously identified. As a result, most visual attention capacity would be devoted to single letter processing in Arabic novice readers, so that reading performance would mainly depend on the successive identification of each letter. In contrast, in the Indo-European languages that use the Latin alphabet, reading fluency would mainly depend on the amount of attention devoted to multi-letter processing, which would determine the size of the orthographic units identified as wholes during serial processing, with a direct impact on reading fluency. Further research is necessary to assess this hypothesis more directly. For example, examining reading performance for the same spoken words written using either Arabic letters or Latin letters (see for example [112]) might help understanding the specific effect of letter complexity on reading fluency. The way visual attention is distributed over the letter string for Arabic word reading might also be directly investigated using specific cueing paradigms (for example, [113]).

## Implication for practice

This study underscores the importance of a comprehensive approach to early literacy instruction that prioritizes both letter recognition training and visual attention skills, especially for Arabic readers. Given the pivotal role of letter knowledge (LK) in reading acquisition, it is essential for children to receive dedicated instruction that focuses on recognizing and naming Arabic letters, as well as understanding their corresponding sounds. This foundational skill is crucial for developing decoding skills and word recognition, which are vital for reading fluency and comprehension. Additionally, the visual complexity of Arabic script highlights the importance of visual attentional span (VAS) in processing letters accurately. Incorporating targeted visual attention exercises, such as visual search tasks [114], or activities that involve simultaneous processing of multiple letters or symbols [115], can aid students in improving visual discrimination skills and attentional focus. Educational curricula should be designed to integrate these insights, ensuring that both letter recognition and visual attention are emphasized in early literacy instruction. By doing so, educators can provide a more comprehensive framework for reading instruction, addressing the specific challenges posed by the Arabic script to beginning readers. Furthermore, for children who are poor readers or have dyslexia, speech and language therapists play a crucial role in screening and intervention. Targeted interventions focusing on the enhancement of letter recognition and visual attention skills [116] should help these children overcome the challenges associated with reading in Arabic. While these practices are important, further research is needed to explore the unique features of the Arabic language and how they impact reading development. Understanding these specifics will enhance our ability to tailor effective interventions and improve literacy outcomes for all learners. The Arabic language’s distinctive characteristics warrant continued attention in research to fully address the diverse needs of readers and to develop more nuanced and effective educational strategies. This holistic approach not only supports typical readers but also equips struggling readers with the necessary skills for lifelong literacy success, thus ensuring that all learners have the opportunity to achieve proficiency in reading.

## Supporting information

**Table S1.** Median (Mdn), Minimum (Min.), Maximum (Max.), Skewness (Skew.) and Kurtosis (Kurt.) after data transformation and normalization for each the variables used in the analysis.

**Table S2.** Results of Linear Regressions with Syllable Reading as the Dependent Variable

**Table S3.** Results of Linear Regressions with Word Reading as the Dependent Variable

**Table S4.** Parameters of the structural equation model to predict syllable reading fluency

**Table S5.** Parameters of the structural equation model to predict word reading fluency

## Credits: Author Contributions

Conceptualization: AG, SV – Data Curation: AG – Formal analysis: ET – Funding: DG, SV – Investigation: AG – Method: AG, SV – Project administration: DG, ET – Supervision: DG, SV – Visualization: ET – Writing (original draft): AG, ET, SV.

## Acknowledgments

We would like to express our sincere gratitude to all the children who participated in our study, as well as their families and school administrators. Without their support, this research would not have been possible. Special thanks to the graduate and undergraduate speech and language therapists who participated in the data collection and data entry processes. Farah, Zeinab I, Abir, Helene, Amira, Banin, Sarah, Zeinab K., Zeinab T., Fatima H, Hanin, Zahraa, Fatima D and Viviane —their help has been invaluable, and their enthusiasm for research and evidence-based practice truly inspiring. We are also grateful to Eric Guinet (LPNC, Grenoble-Alpes University) for his assistance in implementing the Arabic version of the visual attention span tasks. Finally, we extend our thanks to AIRESS and LPNC for their financial support, which made this research possible.

